# Evolution of correlated complexity in the radically different courtship signals of birds-of-paradise

**DOI:** 10.1101/351437

**Authors:** Russell A. Ligon, Christopher D. Diaz, Janelle L. Morano, Jolyon Troscianko, Martin Stevens, Annalyse Moskeland, Timothy G. Laman, Edwin Scholes

## Abstract

Ornaments used in courtship often vary wildly among species, reflecting the evolutionary interplay between mate preference functions and the constraints imposed by natural selection. Consequently, understanding the evolutionary dynamics responsible for ornament diversification has been a longstanding challenge in evolutionary biology. However, comparing radically different ornaments across species, as well as different classes of ornaments within species, is a profound challenge to understanding diversification of sexual signals. Using novel methods and a unique natural history dataset, we explore evolutionary patterns of ornament evolution in a group - the birds-of-paradise - exhibiting dramatic phenotypic diversification widely assumed to be driven by sexual selection. Rather than the tradeoff between ornament types originally envisioned by Darwin and Wallace, we found positive correlations among cross-modal (visual/acoustic) signals indicating functional integration of ornamental traits into a composite unit - the *courtship phenotype.* Furthermore, given the broad theoretical and empirical support for the idea that systemic robustness - functional overlap and interdependency - promotes evolutionary innovation, we posit that birds-of-paradise have radiated extensively through ornamental phenotype space as a consequence of the robustness in the courtship phenotype that we document at a phylogenetic scale. We suggest that the degree of robustness in courtship phenotypes among taxa can provide new insights into the relative influence of sexual and natural selection on phenotypic radiations.

**Author Summary:** Animals frequently vary widely in ornamentation, even among closely related species. Understanding the patterns that underlie this variation is a significant challenge, requiring comparisons among drastically different traits - like comparing apples to oranges. Here, we use novel analytical approaches to quantify variation in ornamental diversity and richness across the wildly divergent birds-of-paradise, a textbook example of how sexual selection can profoundly shape organismal phenotypes. We find that color and acoustic complexity, along with behavior and acoustic complexity, are positively correlated across evolutionary time-scales. Positive covariation among ornament classes suggests that selection is acting on correlated suites of traits - a composite courtship phenotype - and that this integration may be partially responsible for the extreme variation we see in birds-of-paradise.

## INTRODUCTION

Adaptive radiations are driven by ecological differences that promote processes of diversification and speciation [1]. In contrast, phenotypic radiations which occur in the absence of clear ecological differentiation are less well-understood. One commonly investigated mechanism for phenotypic diversification among ecologically similar taxa is variation in social and sexual selection pressures promoting signal or ornament diversification. Ornamental radiations may come about as a consequence of variation in signaling environment [2,3], sensory capabilities [4,5], or pseudo-randomly via mutation order selection [6–8]. Most studies investigating patterns of ornamental diversification have focused on individual trait classes and simplified axes of variation, however, sexual selection does not act on single traits in isolation. A more complete understanding of the processes driving ornamental diversification is possible only by investigating evolutionary relationships between the full suites of ornamental traits under selection.

Many animals rely on multiple ornamental traits to attract mates. Advantages of multiple ornaments may include increased information transfer (multiple messages), increased reliability (redundancy), increased flexibility (ensuring information transfer across contexts and environments), and increased memorability/discriminability [9–13]. Multiple ornaments may be more common when costs associated with the evaluation of those ornaments is low [14], as is likely the case in lekking species [15]. Though we now have broad empirical support for many of the proposed adaptive benefits of multiple signals at the level of individual species, how these specific hypotheses map onto our understanding of phylogenetic patterns of ornament evolution is less clear. Insights into the macroevolutionary implications of multiple ornament evolution are challenging, in part, owing to the difficulties of comparing highly divergent phenotypic traits across species. For example, even focusing on evolutionary patterns of a single trait (e.g. plumage color in birds) across species can be difficult when traits possess different axes of variation (e.g. red vs. blue). Though ingenious new methods have been devised to compare highly divergent ornaments of a single class [16], comparing ornamental complexity across signal classes presents yet an additional layer of complication. However, understanding the interrelationships of different classes of ornaments across phylogenetic scales can potentially provide valuable information about the evolutionary processes of communication, phenotypic radiation, and speciation that cannot be gathered from single trait or single species studies.

If ornamental investment is governed by evolutionary trade-offs, investment in one class of ornaments will come only at the expense of investment in another. Evidence suggests that signal trade-offs manifests as a negative correlation among ornament types across evolutionary time [17–23], reflecting strong, consistent constraints imposed by ecology, physiology, and natural selection [24,25]. Alternatively, instances where ornamental traits show no evolutionary relationships [26–31] suggest long-term patterns of independent evolutionary trajectories or inconsistent selection pressures. In such cases, signals are functionally independent and may even have evolved for use in different contexts (e.g. territorial defense vs. mate attraction). When might we expect positive correlations among ornament classes across species? Theoretical and empirical work on this topic is largely absent, so we propose that positive correlations among signal complexity across species reflect consistent directional selection acting similarly on separate axes of ornamental evolution. Strong, consistent inter-sexual selection could generate these positive correlations, resulting in functional integration among ornament elements [32]. In such cases, positive correlations among signals across species would arise when selection acts upon the ‘integrated whole’ of ornamental traits [33,34], which we call the *courtship phenotype.* The courtship phenotype is the composite expression of all ornamental classes evaluated during courtship and may represent the composite target of selection. Evolution may favor integrated, holistic mate evaluation strategies because of advantages that sensory overlap and redundancy offer (e.g. increased accuracy) [9–13].

Here, we examine broad evolutionary patterns of ornamental signal investment and complexity across the wildly diverse [35,36], monophyletic [37] birds-of-paradise (Paradisaeidae) “…in which the process of sexual selection has gone to fantastic extremes” [38] (Figure 1A). We focus on the birds-of-paradise because this family exhibits extreme variation across species in multiple ornamental axes [35] (e.g. color [39–41], behavior [42,43]), while possessing broadly similar life-histories and mating systems [35,44]. Consequently, insights about the strength, direction, and diversification of ornamental phenotypes in this group may shed light on key processes of sexual selection and its power to generate phenotypic radiation when natural selection-imposed constraints are minimized. In this study, we use a unique natural history dataset on birds-of-paradise to quantitatively evaluate behavioral, acoustic, and colorimetric ornamentation across the 40 species of this family, as well as relationships between signals and display environment.

**Figure 1.**
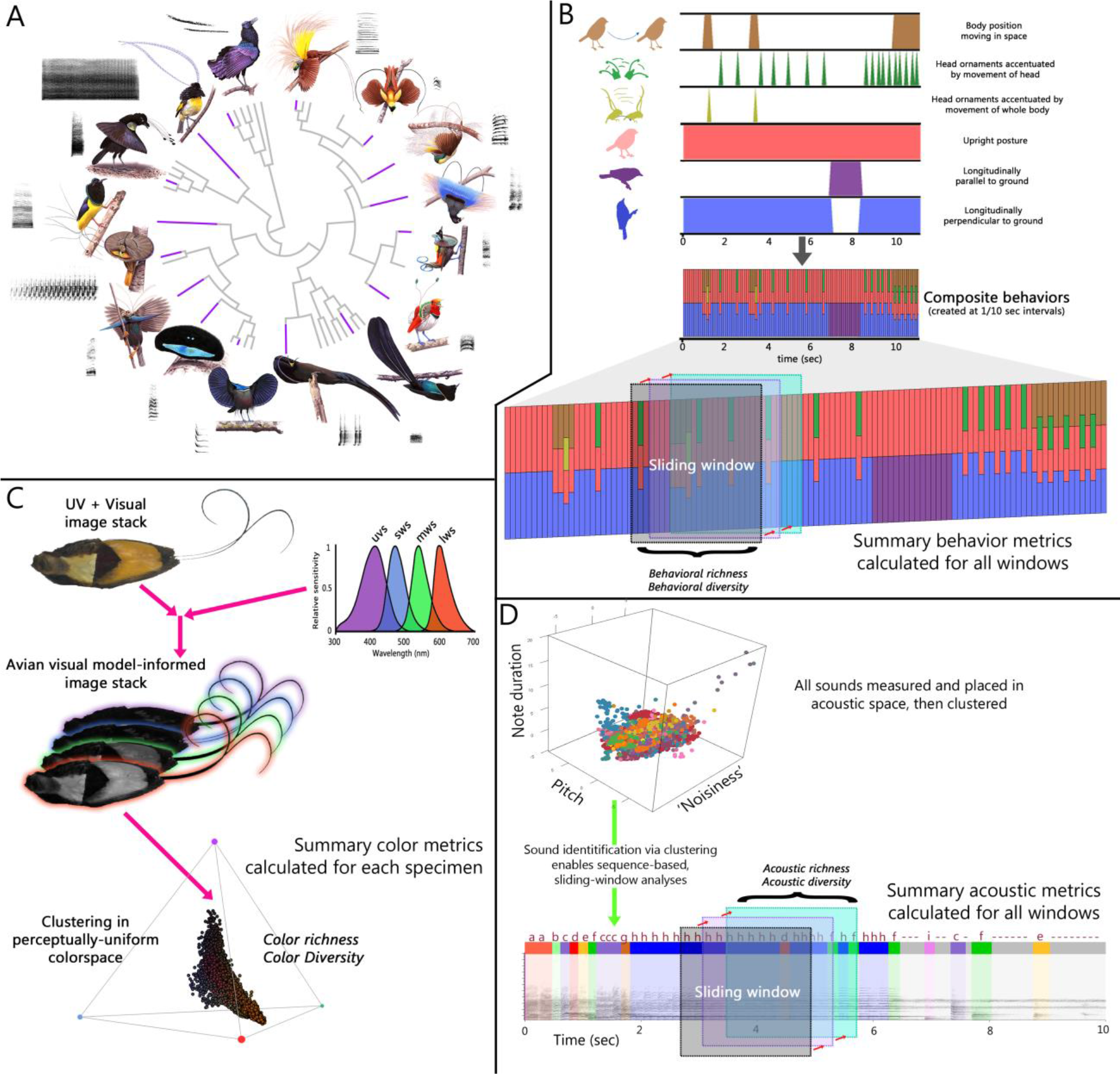
Birds-of-paradise exhibit extreme diversity in colors, sounds, and behaviors used during courtship displays, necessitating novel methods to quantitatively evaluate the evolution of their complexity. (A) Sixteen exemplar species (purple tips) are shown with their phylogenetic relationships to highlight variation in plumage color and courtship display behavior. (B) Behavioral sub-units were scored from field-captured videos of displaying males (Supplementary Tables 3, 4). Behavioral sub-units were combined to create composite behaviors describing any behavior, across species, and facilitating sliding window analysis of behaviors and behavioral sequences. (C) Ultraviolet and visual spectrum images were taken of museum specimens (Supplementary Table 5) and used to generate avian visual model-informed image stacks. Color values were clustered with respect to modeled avian discriminability, enabling whole-specimen quantification of color richness and diversity. (D) All bird-of-paradise sounds were placed into a multidimensional acoustic space defined by principal components analysis. Sounds were then given identities based on locations within acoustic-space, facilitating a sliding-window analysis of sounds and acoustic sequences (Supplementary Table 6, 7).

## RESULTS

### An approach to quantify courtship complexity among divergent ornaments

Comparisons across signal types are inherently challenging for evolutionary biologists given that such signals are necessarily measured in different ways. Additionally, comparisons within color, acoustic, and behavioral repertoires across taxa that vary widely (e.g. the birds-of-paradise) present an additional methodological challenge: how does one compare phenotypes that may share no obvious overlapping characters? We address this obstacle with a two-pronged approach to quantify ornamental complexity for behavior, color, and sounds in the birds-of-paradise. First, we broke down each ornament into a taxonomically-unbounded character space that allowed classification of subunits across all species. Second, we used the specific attributes of a given ornament for each individual, for each species, to categorize the ornament components before quantifying two conceptually-aligned measures of complexity (richness and diversity) for each signal type.

For behavioral analyses, we first broke down the courtship behaviors of all species into distinct sub-units, shared across species (e.g. <u>Supplemental Video 1</u>). We then analyzed composite behavioral sequences across time using sliding-window analyses to compare maximally diverse behavioral repertoires for a set duration across species (Figure 1B). For colorimetric analyses, we relied on visual modeling of multispectral images to quantify the number and relative abundances of perceptually-distinct color types across individuals and species. Though different colors may have different underlying production mechanisms, our analyses simply focused on the number and distribution of distinguishable colors (Figure 1C). Similar to our behavioral analysis pipeline, we used acoustic properties and agglomerative clustering to classify distinct sound-types used by bird-of-paradise in courtship contexts before employing a similar sliding-window analysis to identify maximally diverse acoustic sequences, facilitating comparisons across species (Figure 1D).

In total, we analyzed 961 video clips, 176 audio clips, and 393 museum specimens. From these analyses, we obtained quantitative diversity and richness metrics of ornamental complexity across the birds-of-paradise, which allowed us to rigorously evaluate patterns of correlated character evolution, as well as facilitating our investigation of the influence of breeding system and display environment on ornamental complexity.

### Integrative evolution of courtship complexity across modalities

Using multiple phylogenetic least squares analyses, which allowed us to control for the non-independence of species due to their shared evolutionary history, as well as the potentially confounding influences of display environment (both display height and proximity to courting conspecific males), we uncovered positive correlations between color and acoustic diversity (Figure 2A), as well as between behavioral and acoustic diversity (Figure 2B), suggesting that directional selection has acted similarly on these axes of ornamental complexity. Interestingly, however, there was no significant relationship between color and behavioral diversity, indicating independent evolutionary trajectories for these visually-encoded aspects of courtship ornamentation (Supplementary Table 1). Analyses of ornamental richness revealed similar pattern to those uncovered for ornamental diversity (Supplementary Table 2). Specifically, behavioral and acoustic richness were strongly correlated (as with the corresponding metrics of diversity), while color and acoustic richness were not (in contrast to the pattern for diversity).

**Figure 2.**
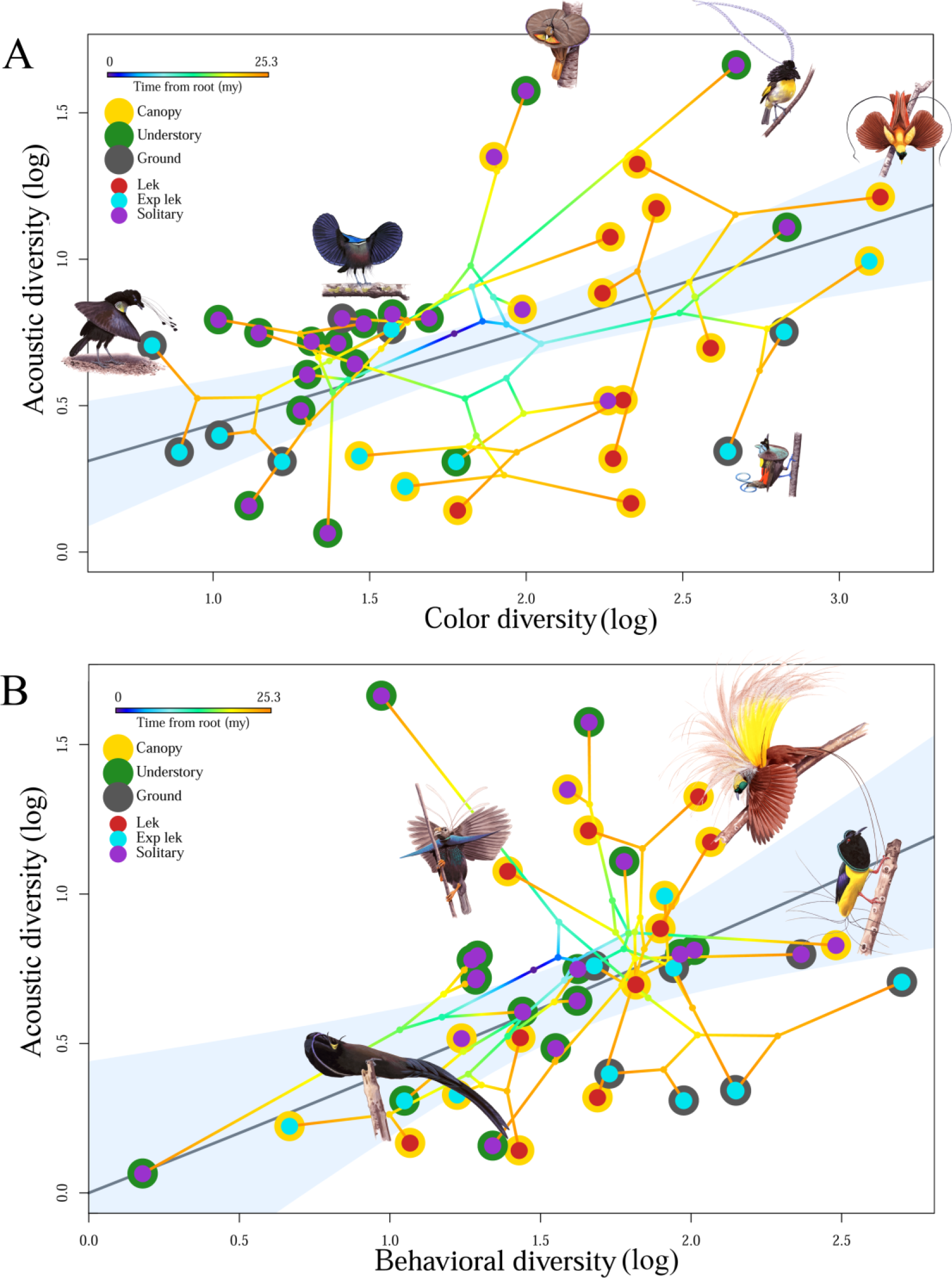
Positive phylogenetic correlations exist among several ornamental diversity indices at an evolutionary scale. (A) Color and acoustic diversity are positively correlated in the birds-of-paradise, where species with greater color diversity exhibit increased acoustic diversity when controlling for behavioral diversity, display height, and display proximity in a multiple phylogenetic least squares regression (mPGLS; summary statistics in Supplementary Table 1). (B) Behavior and acoustic diversity are positively correlated in the birds-of-paradise, where species with greater color diversity exhibit increased acoustic diversity when controlling for behavioral diversity, display height, and display proximity in a multiple phylogenetic least squares regression (mPGLS; summary statistics in Supplementary Table 1).

### Courtship complexity related to spatial distribution of displaying males

All three aspects - behavior, color, sound - of courtship diversity were influenced by the social aspects of display environment (Supplementary Table 1). Lekking species that display in close proximity to one another exhibited more diverse colors and behaviors (Figure 3A, B), corresponding to the increased strength of sexual selection on males to ‘stand out’ visually when being evaluated simultaneously in lekking contexts. Additionally, we found species that displayed solitarily exhibited the greatest acoustic diversity (Figure 3C), perhaps as a means to attract potential mates from greater distances. Color and behavioral richness also increased with display proximity, such that species which display closer together (e.g. in leks) exhibit more colors and behaviors (Supplementary Table 2), and indicating that these patterns are robust to the specific complexity metric used to quantify ornamentation.

**Figure 3.**
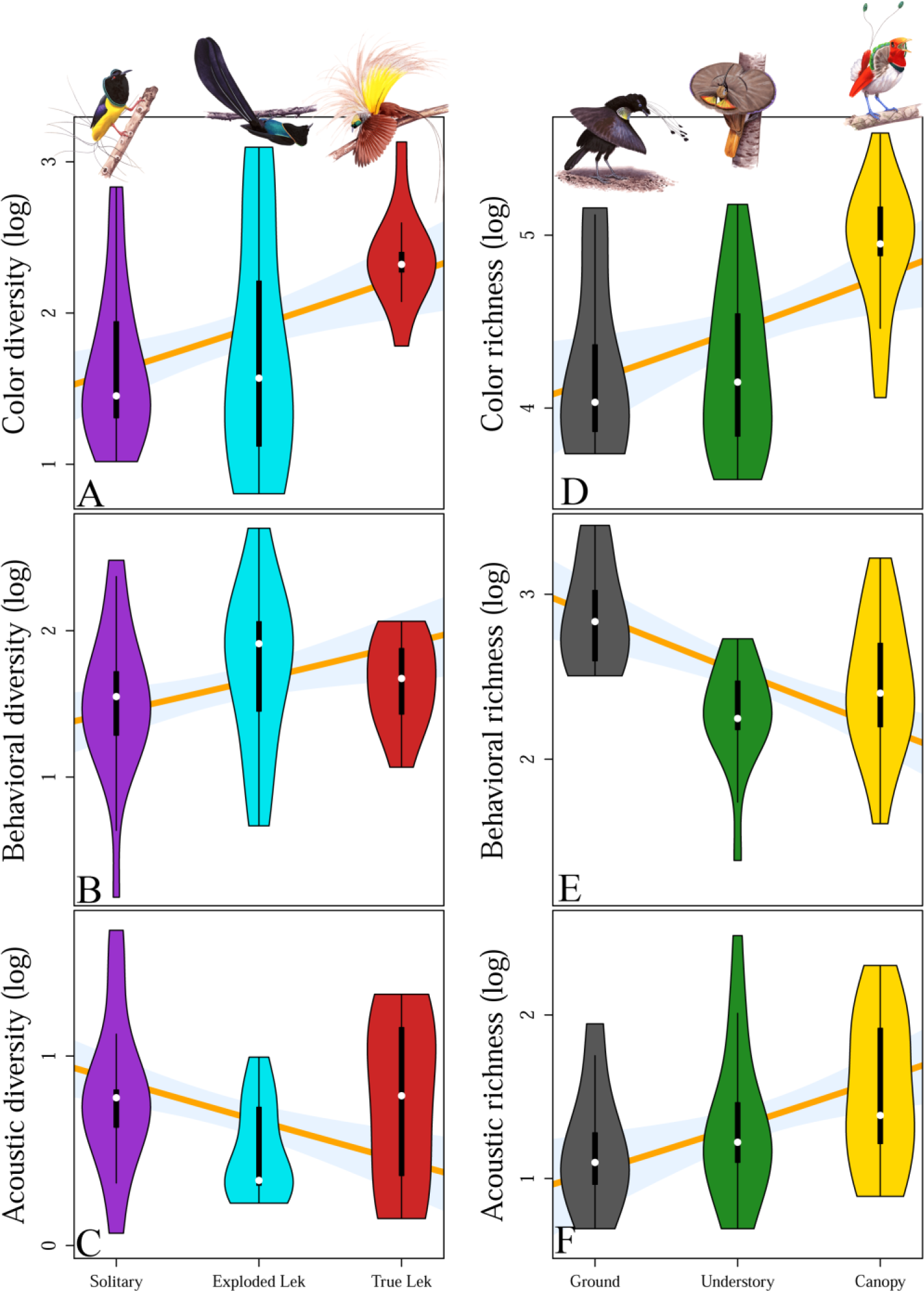
Social and environmental variation in display microhabitat influences multiple axes of ornamental complexity in birds-of-paradise. (A-C) Social dynamics of the display environment, measured as the proximity of other courting males, influences color, behavior, and acoustic diversity in the birds-of-paradise. Species that display in leks have greater color and behavioral diversity (A, B), while species that display solitarily have greater acoustic diversity (C). Additionally, display height influences ornamental richness (D-F), such that species displaying up in the canopy have more colors and sounds (D, F), while species displaying on the forest floor exhibit larger behavioral repertoires (E). Violin plots illustrate the distribution of log transformed diversity (A-C) and richness (d-f) scores for each species, and the orange lines behind the ‘violins’ represent the best-fit lines from multiple phylogenetic least squares (mPGLS) models. Additionally, the shaded regions around each best-fit line represent confidence intervals based on 10^5^ random samples from the multivariate normal distribution of model parameters with the other parameters held at mean empirical values (illustrating the exclusive influence of display proximity (A-C) or display height (D-F) on courtship diversity).

### Courtship complexity related to display height

Among birds-of-paradise, we found that behavioral diversity exhibited a negative relationship with display height, such that species that display down towards the forest floor exhibited the most diverse courtship behavioral sequences (Supplementary Table 1). Species that display in on or near the forest floor are typically operating with lower-light environments, and consequently appear to rely more heavily on complex dance sequences to attract mates. However, neither color nor acoustic diversity were influenced by display height. Interestingly, display height seems to play a more pronounced role in shaping evolutionary patterns of ornamental richness than for ornamental diversity: all richness metrics were influenced by display height (Figure 3D-F). Birds-of-paradise tend to show increased color (Figure 3D) and acoustic (Figure 3F) richness as their display locations increase in height, a result that corresponds to the predictions of sensory drive [45,46] whereby the additional light and openness of the upper-canopy favors increasingly complex color and acoustic displays. Additionally, as with behavioral diversity, behavioral richness was highest for the ground displaying birds-of-paradise (Figure 3E).

## DISCUSSION

Our study provides evidence that selection has favored similar levels of ornamental diversity across multiple signals among the birds-of-paradise. This pattern of positive correlation among distinct ornament classes across evolutionary time-scales and species suggests strong sexual selection on functionally integrated *courtship phenotypes*. The degree to which phenotypic traits are co-expressed and functionally dependent upon one another can be referred to as functional integration [47] or interdependence [48]. Courtship phenotypes with greater functional integration are, therefore, composed of ornaments that are typically expressed at similar levels and which are mutually interdependent in order to influence mate choice [32–34]. Correlations among the signals that comprise the courtship phenotype also suggest a previously undescribed robustness in bird-of-paradise courtship phenotypes that may have played a key role in the extreme ornamental radiation exhibited by this taxon (Figure 1).

Evolutionary biologists dating back to Mayr [49] and even Darwin [50] have recognized the potential evolutionary implications of functional redundancy (where two or more structures perform the same function). Redundancy facilitates evolutionary innovation due to the “backup” nature of systemic redundancy (e.g. this is a key idea regarding the evolutionary benefit of diploidy [51,52]). The term ‘robust’ describes systems where the overall structure and interconnectedness of parts provides protection from environmental or mutational instability [53], where a given function is not lost if a key component fails - not due to a duplicated structure but rather piecemeal interdependencies. All redundancy provides a measure of robustness, but robust systems are not necessarily redundant[54]. Given the broad theoretical [55–57] and empirical [58–60] support for the idea that robustness can promote evolvability across a wide array of biological domains, we posit that the correlations among signal types within birds-of-paradise courtship phenotypes are, at least partially, responsible for the dramatic diversification and radiation of courtship signals displayed by birds-of-paradise. If female birds-of-paradise make mate choice decisions based on sensory input from the multiple signals that comprise a composite courtship phenotype, and information from those channels is correlated, then novel mutations changing the structure or form of a given ornament may occur without “necessary” information being lost [61]. Consequently, over evolutionary time, the robustness of courtship phenotypes can also lead to their collective diversification as a by-product of increased evolvability.

Phenotypic radiations in the absence of clear ecological differentiation may arise stochastically [6,7] and be heavily influenced by the specific intricacies of female choice [8,62–64]. Birds-of-paradise clearly exhibit some ecological differentiation [35] but, broadly speaking, they tend to be heavily frugivorous and predominantly polygynous [65]. They do not, however, all display to potential mates in the same contexts or microenvironments. Some species display high in the canopy, some down on the forest floor, and others in the understory in between. Likewise, some species display in large, cacophonous leks, some in exploded leks where males can hear but not see one another, and other species display solitarily. Our results suggest that these differences have shaped the specific courtship and signaling strategies of each species (Figure 3). More colorful birds display high in the canopy where abundant light increases the likelihood that females will be able to detect and discern numerous, elaborate colors. More behaviorally complex birds tend to display near the forest floor where there is less light (and ability to perceive subtle variation in color) but more area available for a courtship stage or “dance floor”. Birds that display solitarily have more diverse acoustic repertoires, perhaps because females can identify attractively signing individuals (their voices will not get lost in a crowd). Display-site and display-context thus influence the specific forms of ornamentation possessed by individual species, and taking them into account from an analytical perspective allows us to better understand patterns of signal coevolution and the potential importance of a functionally integrated courtship phenotype.

Signal efficacy and information content can exert strong influence on receiver preferences, and understanding both elements is integral when examining the evolution of complex, multicomponent courtship phenotypes [11,66,67]. For example, the perceptual abilities [68,69] and psychology of signal receivers [70,71], as well as the environments through which signals are transmitted [45], can markedly influence signal efficacy. Additionally, the information content of multiple signals may increase the net amount of information transferred (e.g. multiple messages [13]) or increase accuracy and reliability if multiple signals communicate the same message (e.g. redundant signals [9,13]). The perceptual channels by which birds-of-paradise attract mates and those channels that are correlated at a phylogenetic scale provide tantalizing, though tentative, insights into the processes of efficient information transfer and receiver stimulation regulating mate choice in this group. Specifically, the fact that significant positive correlations exist between acoustic and color signals (auditory, visual), and between acoustic and behavioral signals (auditory, visual), but *not* between color and behavioral signals (visual, visual) aligns with psychometric literature on information and sensory input. When multiple sources of information are provided, information may be maximized if that information comes from separate channels (e.g. acoustic, visual) and lost when arriving through a single sensory channel [72] (but see [73]). What exactly this ‘information’ might be in birds-of-paradise (quality [74], attractiveness [62], motivation [75], etc.) is not clear, but this result provides an interesting starting point for future investigations.

Phylogenetic comparative investigations of animal signals hold the potential to answer important questions about the evolutionary trajectories of communication over time [76,77]. However, the accuracy of the insights obtained from such investigations depend on the (i) quality and (ii) completeness of the data analyzed. First, there exist well-established, validated data collection protocols that allow researchers to make rigorous empirical comparisons among taxa for many animal signals (e.g. color, sound). To ensure the validity of the conclusions drawn from comparative studies of animal signals, these approaches need to be implemented. As an example, a recent study [78] using human-scored sexual dichromatism scores based on color paintings revealed no support for the idea that sensory drive [45] influences the plumage coloration of male birds-of-paradise. In contrast, the results we obtained using high-definition, full-spectrum photography integrating avian visual modeling suggests that sensory drive has influenced color and acoustic complexity in the birds-of-paradise. Furthermore, behavioral classification obtained from written descriptions from multiple sources is insufficient to provide meaningful insights into actual behavioral sequences or the relative temporal investment of individuals in the various behavioral states. Though we still may be a way off from automated, high-throughput behavioral classification via machine learning (*sensu* [79]) in natural contexts, there exist repositories and collections of vouchered video data from diverse taxa enabling the consistent, high-fidelity, interpretable quantification of behavior [80] necessary for in-depth insight into behavioral evolution. Second, partial datasets used to investigate evolutionary interdependencies can be misleading. If researchers are interested in how a single class of ornaments has changed over evolutionary time and are interested in environmental characteristics associated with those changes, analyzing a single ornament is a logical approach. If, however, researchers are interested in the evolutionary relationships among signals across phylogenetic time-scales [30], then the inclusion of only some ornamental characters can lead to erroneous conclusions. For example, it is not uncommon to focus only on color and behavior displays [21,78] (i.e. ignoring the role that acoustic signals can play in complex courtship displays), or only on color and acoustic displays [20,26] (i.e. ignoring behavioral displays). In some cases, such decisions make good sense (as in groups lacking acoustic elements or complex behavioral displays), but when datasets omit key signal categories erroneous conclusions can be reached (e.g. those outlined in [78], focused only on color and behavior). Indeed, one our key findings - that positive correlations exist among different axes of courtship phenotypes at evolutionary scale, suggesting selection for overall signal robustness - was made only because we included color, behavior, and sound in our analysis. In many respects, these issues are not specific to evolutionary studies of animal signals (i.e. low-resolution or incomplete data are never ideal), yet we highlight them here in part owing to the rapidity and complexity of the developing methodologies used to study animal communication and to illustrate the fact that upstream data acquisition decisions can have significant consequences on our empirical interpretations of evolutionary processes.

Evolutionary trade-offs - increases in trait expression linked to reductions in another - are ubiquitous: “If there were no trade-offs, then selection would drive all traits correlated with fitness to limits imposed by history and design” [81]. Hence, the finding that the ornaments of birds-of-paradise are positively correlated at phylogenetic scale suggests that the evolution of these traits occurs relatively unhindered by natural selection, ecology, or physiological limits. Consequently, sexual selection is the most likely candidate as a driver of phenotypic radiation in birds-of-paradise. The correlation among ornamental classes also suggests that selection is acting on functionally integrated *courtship phenotypes* for birds-of-paradise, a finding that suggests female birds-of-paradise make mate choice decisions incorporating holistic, multicomponent information sets comprised of the various ornaments possessed by males of their species. Rather than being unique to birds-of-paradise, however, we suggest that this phenomenon is widespread among animals - though is at varying degrees constrained, impeded, or obfuscated by conflicting and constraining processes and limitations imposed by ecology and natural selection. The degree to which selection has facilitated the evolution of integrated, robust *courtship phenotypes* may in fact serve as a proxy for the overall strength and consistency of female-driven sexual selection in any taxa, where the integration and correlation among ornaments comprising the courtship phenotype may shed important light on the history and strength of sexual selection in that particular group.

## METHODS

### Behavioral complexity

We quantified the behavioral complexity of courtship display behaviors for the birds-of-paradise by scoring field-recorded video clips of 32 (80%) paradisaeid species, primarily from the Macaulay Library at the Cornell Lab of Ornithology (macaulaylibrary.org, Supplementary Table 3). In total, we watched 961 clips from 122 individuals totaling 47707.2 seconds (≈795.12 minutes; mean clip duration = 49.64s). Courtship display behavior is highly variable among BOP species, necessitating broad behavioral categories to facilitate investigations of behavioral evolution. Specifically, one of us (CDD) blindly evaluated video clips of male BOPs displaying species-typical courtship behaviors [35] using a customized ethogram of behavioral units that enabled us to quantify all state and event behaviors exhibited by all species of Paradisaeidae (Supplementary Table 4).

#### Data collection

To record courtship display behaviors, we used a customized version of an open-source behavior logging program [82]. Additionally, we created a customized keyboard that allowed us to quickly and accurately record the start/stop times of all duration behaviors, as well as the instances of all event behaviors. The combinations of different behavioral categories throughout each clip allowed us to generate sequence data of distinct behavioral elements.

#### Measures of sexual display behavior complexity

Courtship displays can be broken down into distinct behavioral elements and the transitions between these elements. We investigated the number of unique behavioral elements (*behavioral richness*) in a given time period, as well as the Shannon entropy [83] of these behaviors (*behavioral diversity*). Shannon entropy provides a measure of ‘information’ encoded in the behavioral displays, and we converted Shannon entropy scores to their numbers equivalents [84,85]. Shannon indices were chosen specifically because they are the only measures that “give meaningful results when community weights are unequal” [84]. As previously described [84], the numbers equivalent for Shannon entropy values has the readily interpretable property whereby a value of 2x would indicate a behavioral sequence with twice as many equally-well-represented behaviors as a sequence with a value of x.

#### Sliding window analysis

The number of courtship recordings available was highly variable across species of BOPs (Supplementary Table 3). To reduce the influence of sampling intensity on our overall behavioral analyses, we used a sliding-window analysis to evaluate similar time windows for courtship display complexity across species. Specifically, we used a sliding 50s window across all clips for a given individual to identify the specific 50s period of maximal display complexity for that individual, and incorporated only the resultant complexity scores for this interval in our analysis. This approach minimizes the influence that variation in recording time and clip duration has on species-level behavioral comparisons. Our results and interpretations are robust to the choice of different window sizes between 20 and 60 seconds (Supplementary Figure 1,2).

### Color complexity

#### Image collection

We collected images from 393 BOP museum specimens (Supplementary Table 5) housed at the American Museum of Natural History. Specifically, we took RAW format images of adult males from 40 BOP species under standardized conditions using a Canon 7D camera with full-spectrum quartz conversion and fitted with a Novoflex Noflexar 35 mm lens. Illumination was provided from two eyeColor arc lamps (Iwasaki: Tokyo, Japan) diffused through polytetrafluoroethylene sheets 0.5mm thick. These arc lamps are designed to simulate CIE (International Commission on Illumination) recommended daylight (D65) illumination (though they come standard with a UV-blocking coating, which we removed prior to use). Additionally, for every specimen position (see below) we used filters (Baader: Mammendorf, Germany) to take two photos, one capturing only ultraviolet light (300-400nm) and one capturing wavelengths between 400 and 700 nm.

To simulate a variety of viewing angles and increase the likelihood of capturing relevant coloration from bird specimens, we took photographs of each specimen from three viewing angles: dorsal, ventral, ventral-angled. Specifically, each specimen was photographed from above while it was flat on its belly (dorsal view), flat on its back (ventral view), and angled 45° on its back (rotating the frontal plane along the vertical-axis, while keeping the head oriented in the same direction as the previous two photographs). The angled photograph was taken to increase the likelihood of capturing some of the variation made possible by iridescent plumage.

#### Image processing

Ultra-violet and visible spectrum images were used to create standardized (i.e. channels were equalized and linearized [86]) multispectral image files for each specimen/position using the Image Calibration and Analysis Toolbox [87] in ImageJ [88].

##### a. Avian color vision

After estimating the color sensitivity of our camera/lens combination [86,87], we generated custom mapping functions to convert image colors to stimulation values corresponding to avian cones. The spectral sensitivities of BOPs are not currently known, so we evaluated color using a visual model from another passerine species (the blue-tit [89]). Prior to subsequent analysis, we performed a median pixel blur to eliminate aberrant pixel values (owing to dust on the sensor, temporary dead-pixels, etc.).

##### b. Color clustering

Following conversion to avian color vision and noise filtering, we used agglomerative hierarchical clustering to reduce each multispectral image down to a perceptually-relevant number of color clusters. At the first step of the clustering process, each pixel is its own cluster. Each cluster is then compared to its neighboring clusters in the XY plane of the image, and composite distances calculated based on an equal weighting of chromatic [90] and achromatic [91] Just Noticeable Distances. Following distance calculations, each cluster is combined with its nearest neighboring cluster. Nodes can have multiple clustering events at each pass, e.g. if cluster A is closest to B, but B is closest to C then all three will be clustered, meaning whole strings or neighboring regions can be clustered. In practice, each cluster tends to be combined with two or three clusters on each pass. Additionally, the distance radius increases with each pass, so as clusters get larger they can also combine with neighboring clusters further away which has the desirable effect of also keeping the processing for each pass relatively constant (i.e. there are fewer clusters on each pass, but each one must be compared to a larger number of neighbors). Clustering therefore takes place across n + 2 dimensions (n colors plus x and y space).

##### c- Measures of color complexity

Following color clustering, we quantified plumage color complexity using analogous indices to those we employed in our behavioral analysis. Namely, we quantified color richness (the number of distinct clusters and color diversity (the numbers equivalent of Shannon index).

##### d- Influence of specimen age on color complexity measures

Aging can influence the coloration and appearance of some kinds of avian plumage [92,93], though such effects are often relatively small [94]. To evaluate the possibility that specimen age might influence our estimates of species’ level plumage elaboration, we conducted a linear mixed-effect model with two measures of color complexity (color richness, color diversity) as the dependent variable, collection year as the independent variable, and species as a random effect. Analyzing these models revealed no significant influence of collection year on either color richness (standardized *β* = 0.032, 95% CI −0.036 - 0.099, *t* = 0. 930, *p* = 0.353) or diversity (standardized *β* = 0.033, 95% CI −0.012 - 0.080, *t* = 1.437, *p* = 0.151).

### Acoustic Complexity

As with display behaviors, we quantified the acoustic complexity of courtship sounds produced by analyzing field-recorded audio/video clips of 32 (80%) BOP species. In total, we analyzed sound from 176 clips from 59 individuals totaling 24670.9 seconds (≈411 minutes; mean clip duration = 140.18s; Supplementary Table 6). Though birds can generate sounds (both vocally and mechanically) in numerous contexts, we focused our analysis on recordings from known display sites or those matching written descriptions of courtship sound production[35].

#### Data collection

From each video clip used to quantify display behavior we identified a focal individual and all of the sounds it produced. Spectrograms of the audio were viewed with a frequency resolution of 43.1 Hz and time resolution of 2.31 ms, and all sounds were marked in the sound analysis software RavenPro v. 1. 5[95]. Individual sounds were defined as temporally-separated sound elements. Using the robust measurements in Raven, we measured the duration, maximum and minimum frequency, bandwidth, peak frequency, and peak frequency contour of each call. We measured the disorder in a call with aggregate entropy and average entropy measures in Raven.

Following detailed analysis of the acoustic parameters for all notes, we used a two-step semi-automated classification analysis to assign note identity. In the first step, we conducted a principal components analysis using 15 summary acoustic variable (Supplementary Table 7) followed by agglomerative hierarchical clustering to assign partial note identity. In the second step, each note was given a categorical identifier depending on the combination of four qualitative variables manually scored as yes/no (frequency modulation, non-harmonic structure, impulsive, stochastic). Full note identity was achieved by merging the clustered note identity with the combined qualitative-categorization.

#### Measures of acoustic complexity

After assigning identities to all notes in our dataset, we measured acoustic richness (number of distinct note types) and acoustic diversity (Shannon index of notes) within a given time period (see *Sliding window analysis* below). As with behavior, we used the numbers equivalent of Shannon index values to facilitate more direct comparisons among samples and species.

#### Sliding window analysis

The duration and number of available courtship-specific acoustic recordings was highly variable across species of BOPs (Supplementary Table 6). To reduce the influence of this variation on species-level acoustic comparisons, we used a sliding-window analysis, similar to our behavioral analyses, to evaluate and compare similar time windows for acoustic display complexity across species. To identify the time period of maximal acoustic complexity for an individual in our analysis, we used a sliding 10s window, chosen as the minimum duration resulting in relatively stable individual complexity scores (Supplementary Figure 3), across all clips for a given individual. Relative complexity measures are robust to the choice of different window sizes between 5 and 50 seconds (Supplementary Figure 4, 5).

### Phylogenetic analyses

#### Tree

We regenerated a phylogenetic hypothesis from the nexus format alignment provided to us by the authors of a recent molecular phylogeny for Paradisaeidae [96]. We applied the same partitioning and substitution model scheme as described in the original publication, and our re-analysis of the original data set generated an identical phylogeny, with branch lengths proportional to substitutions/site. We then estimated an ultrametric time-scaled tree for downstream comparative analyses with the semi-parametric rate smoothing algorithm [97] implemented in the *chronopl* function in the ape v4.1 R package [98]. We set the root age to 50 million years [99] to approximate the age of the split between Acanthasitta and other Passeriformes, and scaled the remaining branches accordingly. Additionally, we modified our tree topology to accommodate the revised taxonomic relationship of *Lophorina superba* [100], placing this species as the outgroup to *Ptiloris*.

#### Phylogenetic Generalized Least Squares

For each of the six components of courtship phenotype (color richness and diversity, behavior richness and diversity, acoustic richness and diversity), we conducted a single, multiple phylogenetic least squares (mPGLS) regression evaluating the influence of the other elements of courtship phenotype and two signal-environment variables predicted to influence relative investment in separate axes of overall courtship phenotype. Specifically, we included a pseudo-continuous metric of display height in all models, where each species was scored as displaying on the forest floor, in the understory, or in the forest canopy. Additionally, we included the pseudo-continuous metric of display proximity in all models, where each species was scored as displaying solitarily, in exploded leks, or in true leks. In each model (see Supplemental Table 1, 2), we included only ‘like’ phenotype measures (e.g. including behavioral and acoustic richness, but not behavioral or acoustic diversity, when investigating the drivers of color richness). All courtship phenotype measures were log-transformed prior to analyses.

#### Imputation

We used the Rphylopars package in R [101] to impute character values for taxa with missing data (e.g. species lacking behavioral/acoustic information, Supplementary Table 4). This methodology has previously been found to perform well in predicting ancestral and missing species’ values [102]. In our case, we evaluated the performance of several methods to estimate missing values assuming i) a Brownian motion model of trait evolution, ii) an Ornstein-Ulbeck model[103], iii) an “early-burst” model of trait evolution [104], iv) a Pagel’s lambda model of trait evolution [105], and v) a multivariate OU model [106]. We compared model performance by evaluating AIC scores and determined that the OU model performed best. Consequently, character trait values imputed using this model were used in all subsequent analysis.

Though data imputation can increase statistical power [107], the instances in which it might induce spurious findings are few (especially given the relatively small proportion of our total dataset (20%) for which we imputed values (*cf* [108,109]). In fact, bias tends to be lower when missing data are imputed rather than omitted [102]. Regardless, to alleviate concerns that imputed values may drive subsequent findings, we also conducted our phylogenetic least squares (PGLS) analyses on the limited subset of species (n=32) for which we have complete data. In all cases, the findings were qualitatively identical to those reported in the main text (Supplementary Table 8, 9).

### Data accessibility

Data will be made available to readers.

## Acknowledgements

We thank Brett Benz, Lydia Garetano, Paul Sweet, and Joel Cracraft at the American Museum of Natural History for their generous help. We thank the Cornell NBB “LunchBunch” for stimulating intellectual contributions. We thank Nick Mason, Jacob Berv, Eliot Miller, Mike Sheehan, Gavin Leighton, Eric Goolsby, Anastasia Dalziell, Marcelo Araya Salas, Leo Campagna, Sara Miller, Irby Lovette, and Mike Webster for input and advice on data collection, analysis, and interpretation. We thank Szabolcs Kókay for the incredible paintings illustrating bird-of-paradise display behavior in Figure 1. We thank Rory, Teddy, Veronica, Sandy, and David Ligon for emotional support. RAL was supported by NSF Grant #1523895.

## Supplementary Material

**Supplementary Video 1: Bird-of-paradise behavioral scoring demonstration. In this video, a male western parotia *Parotia sefilata* performs a species-typical courtship dance for females perching above.** This video demonstrates the concordance between a sub-sample of our scored behaviors and the actual performance of the bird in real-time. The behaviors represent body-position moving (BP1), changing direction while moving (BP2), shape-shifting (SS1), bowing (O3), ornamental head plumage accentuation by moving those feathers (OPMH), ornamental flank plumage accentuation by moving those feathers (OPMF), ornamental head plumage by moving the head (OPAH), and ornamental flank plumage accentuation by moving the torso (OPAC1).

**Supplementary Table 1.**
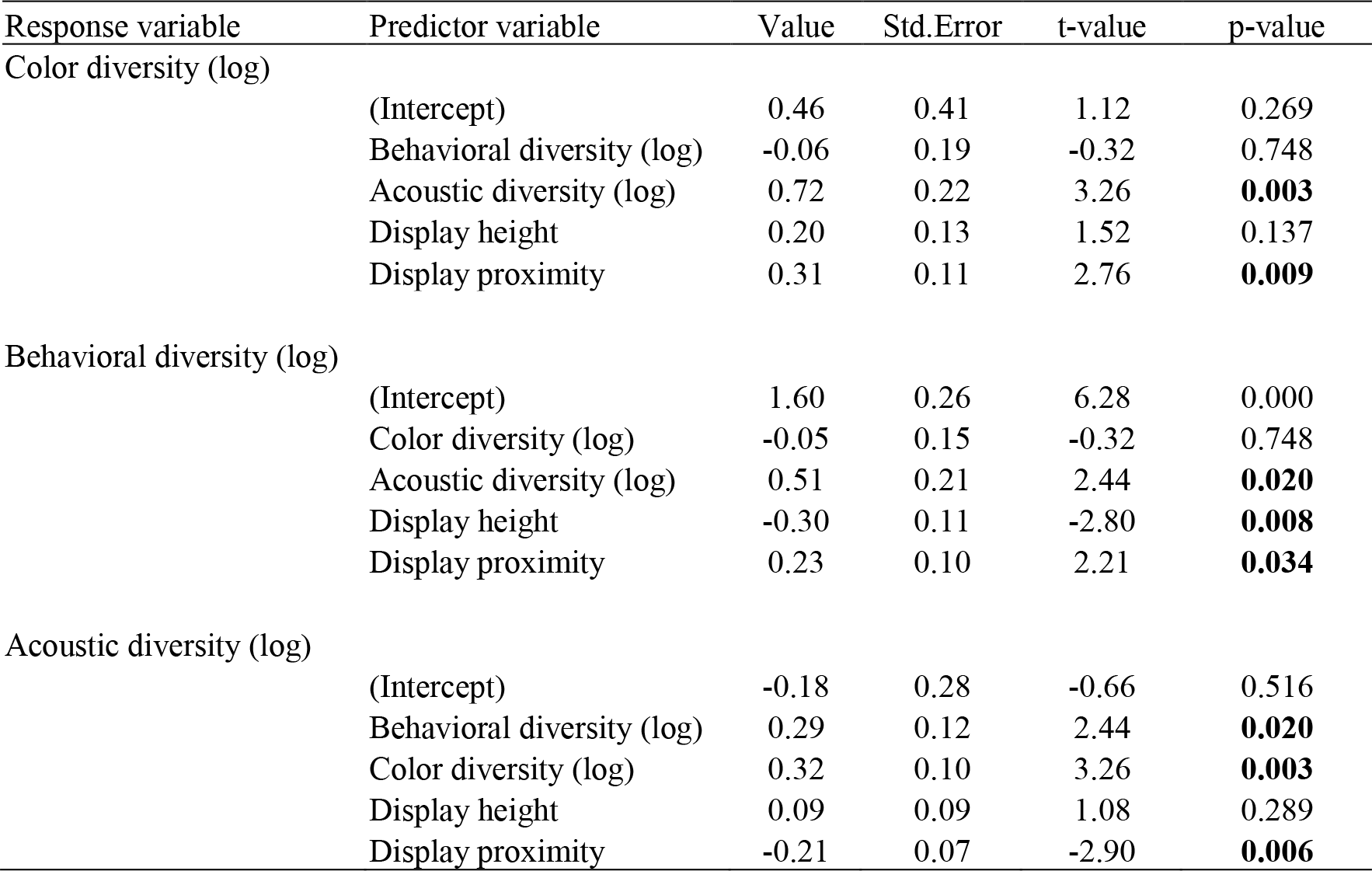
Multiple phylogenetic least-squares (mPGLS) analyses of communication relevant influences on three axes of courtship phenotype diversity.

**Supplementary Table 2.** Multiple phylogenetic least-squares (mPGLS) analyses of communication relevant influences on three axes of courtship phenotype richness.

| Response variable | Predictor variable | Value | Std.Error | t-value | p-value |
| --- | --- | --- | --- | --- | --- |
| Color richness (log) | (Intercept) | 2.91 | 0.57 | 5.15 | 0.000 |
|  | Behavioral richness (log) | 0.13 | 0.21 | 0.62 | 0.539 |
|  | Acoustic richness (log) | 0.15 | 0.18 | 0.81 | 0.425 |
|  | Display height | 0.30 | 0.13 | 2.34 | **0.025** |
|  | Display proximity | 0.23 | 0.10 | 2.19 | **0.035** |
| Behavioral richness (log) | (Intercept) | 2.01 | 0.48 | 4.17 | 0.000 |
|  | Color richness (log) | 0.08 | 0.13 | 0.62 | 0.539 |
|  | Acoustic richness (log) | 0.37 | 0.13 | 2.83 | **0.008** |
|  | Display height | -0.34 | 0.09 | -3.65 | **0.001** |
|  | Display proximity | 0.18 | 0.08 | 2.15 | **0.038** |
| Acoustic richness (log) | (Intercept) | -0.72 | 0.67 | -1.08 | 0.286 |
|  | Behavioral richness (log) | 0.50 | 0.18 | 2.83 | **0.008** |
|  | Color richness (log) | 0.12 | 0.15 | 0.81 | 0.425 |
|  | Display height | 0.28 | 0.12 | 2.45 | **0.020** |
|  | Display proximity | -0.17 | 0.10 | -1.78 | 0.084 |

**SSupplementary Table 3:** Species sampled for courtship behavior, including the number of individuals watched.

| Species | N<br>individuals | N<br>clips | Mean duration<br>of clips (s) | Total time<br>watched (s) |
| --- | --- | --- | --- | --- |
| *Paradisaea raggiana* | 5 | 39 | 24.4 | 950 |
| *Paradisaea apoda* | 2 | 32 | 74.8 | 2392.8 |
| *Paradisaea minor* | 7 | 38 | 44.9 | 1706.3 |
| *Paradisaea rubra* | 7 | 76 | 21.9 | 1663.8 |
| *Paradisaea decora* | 4 | 40 | 56.5 | 2261.2 |
| *Paradisaea guilielmi* | 3 | 9 | 41.4 | 373.0 |
| *Paradisaea rudolphi* | 4 | 22 | 60.5 | 1330.8 |
| *Cicinnurus respublica* | 3 | 39 | 50.6 | 1973.1 |
| *Cicinnurus magnificus* | 4 | 30 | 73.3 | 2197.6 |
| *Cicinnurus regius* | 4 | 29 | 33.3 | 964.4 |
| *Astrapia mayeri* | 5 | 13 | 42.7 | 554.8 |
| *Astrapia rothschildi* | 3 | 20 | 69.0 | 1379.9 |
| *Astrapia splendidissima* | 1 | 4 | 74.1 | 296.4 |
| *Astrapia nigra* | 1 | 4 | 52.4 | 209.6 |
| *Epimachus meyeri* | 3 | 12 | 106.9 | 1282.5 |
| *Epimachus fastosus* | 5 | 25 | 223.3 | 5581.8 |
| *Ptiloris magnificus* | 3 | 20 | 152.1 | 3041.9 |
| *Lophorina superba* | 2 | 8 | 171.2 | 1369.8 |
| *Ptiloris victoriae* | 5 | 8 | 215.0 | 1719.7 |
| *Ptiloris paradiseus* | 5 | 21 | 35.4 | 742.8 |
| *Semioptera wallacii* | 6 | 60 | 15.3 | 917.5 |
| *Drepanornis bruijnii* | 2 | 3 | 69.7 | 209.1 |
| *Drepanornis albertisi* | 1 | 8 | 114.4 | 914.9 |
| *Seleucidis melanoleucus* | 1 | 19 | 48.2 | 915.3 |
| *Parotia wahnesi* | 6 | 174 | 24.7 | 4300.5 |
| *Parotia sefilata* | 2 | 54 | 44.4 | 2396.4 |
| *Parotia lawesii* | 7 | 51 | 34.3 | 1747.4 |
| *Parotia helenae* | 4 | 21 | 32.9 | 691.7 |
| *Parotia carolae* | 12 | 54 | 50.3 | 2714.5 |
| *Pteridophora alberti* | 3 | 24 | 31.8 | 763.3 |
| *Manucodia comrii* | 1 | 3 | 17.2 | 51.5 |
| *Phonygammus keraudrenii* | 1 | 1 | 92.9 | 92.9 |

**Supplementary Table 4.**
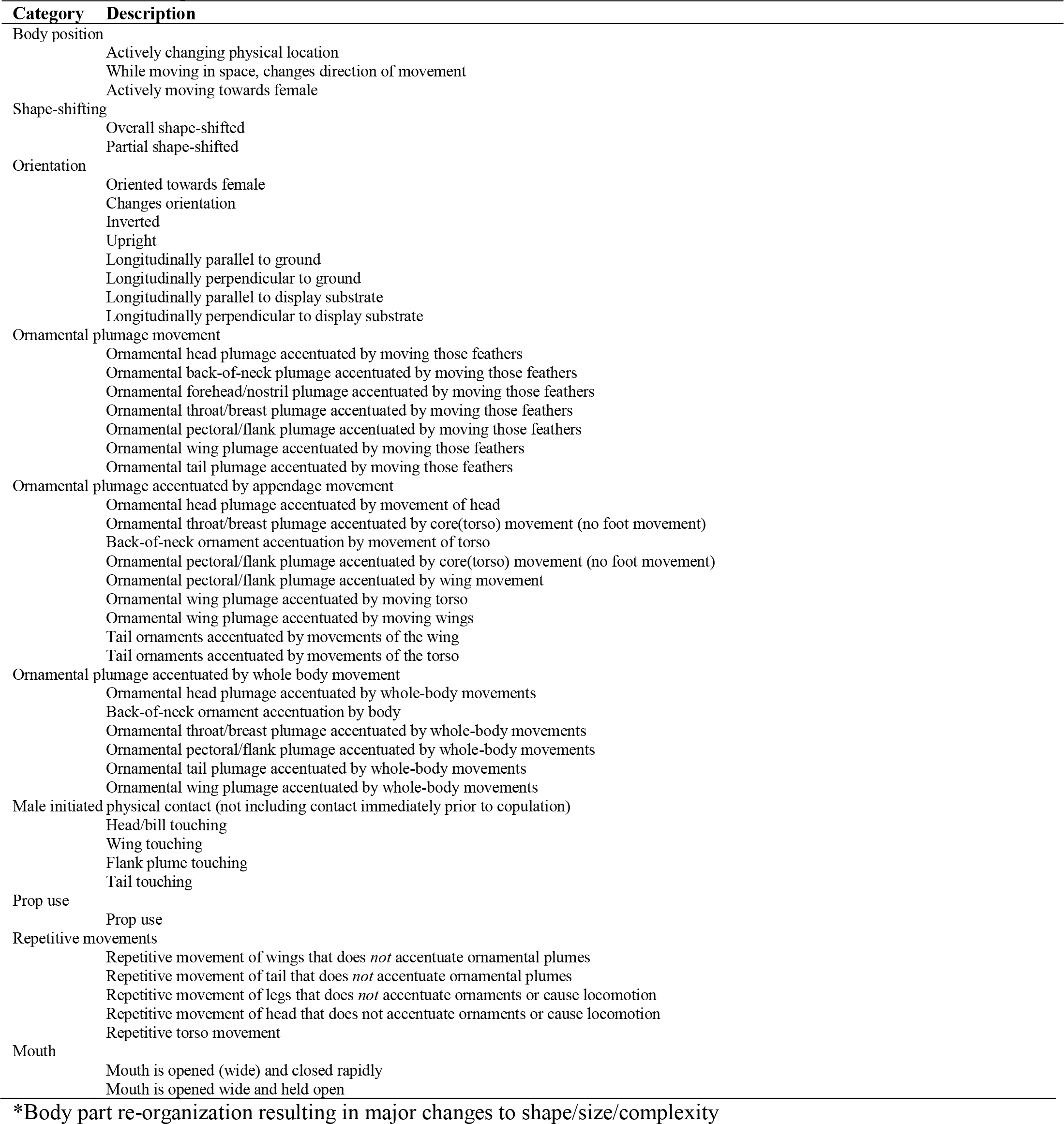
Ethogram describing homologous behaviors scored while observing courtship display behavior of birds-of-paradise.

**Supplementary Table 5.** Summary of specimens used to quantify color complexity in the birds-of-paradise.

| Species | N sampled |
| --- | --- |
| *Paradisaea raggiana* | 10 |
| *Paradisaea apoda* | 10 |
| *Paradisaea minor* | 10 |
| *Paradisaea rubra* | 10 |
| *Paradisaea decora* | 12 |
| *Paradisaea guilielmi* | 10 |
| *Paradisaea rudolphi* | 11 |
| *Cicinnurus respublica* | 10 |
| *Cicinnurus magnificus* | 11 |
| *Cicinnurus regius* | 11 |
| *Astrapia stephaniae* | 10 |
| *Astrapia mayeri* | 10 |
| *Astrapia rothschildi* | 10 |
| *Astrapia splendidissima* | 10 |
| *Astrapia nigra* | 10 |
| *Paradigalla carunculata* | 5 |
| *Paradigalla brevicauda* | 10 |
| *Epimachus meyeri* | 10 |
| *Epimachus fastosus* | 10 |
| *Ptiloris magnificus* | 10 |
| *Ptiloris intercedens* | 10 |
| *Lophorina superba* | 10 |
| *Ptiloris victoriae* | 10 |
| *Ptiloris paradiseus* | 10 |
| *Semioptera wallacii* | 10 |
| *Drepanornis bruijnii* | 13 |
| *Drepanornis albertisi* | 11 |
| *Seleucidis melanoleucus* | 10 |
| *Parotia wahnesi* | 4 |
| *Parotia sefilata* | 10 |
| *Parotia lawesii* | 10 |
| *Parotia helenae* | 3 |
| *Parotia carolae* | 10 |
| *Pteridophora alberti* | 10 |
| *Manucodia comrii* | 10 |
| *Manucodia chalybatus* | 10 |
| *Manucodia jobiensis* | 10 |
| *Manucodia ater* | 10 |
| *Phonygammus keraudrenii* | 10 |
| *Lycocorax pyrrhopterus* | 12 |

**Supplementary Table 6.** Species sampled for acoustic courtship complexity, including the number of individuals watched.

| Species | N individuals | N clips | Mean duration of clips (s) | Total time analyzed (s) |
| --- | --- | --- | --- | --- |
| *Paradisaea raggiana* | 1 | 3 | 94.8 | 284.3 |
| *Paradisaea apoda* | 1 | 8 | 68.0 | 543.8 |
| *Paradisaea minor* | 2 | 4 | 37.5 | 150.2 |
| *Paradisaea rubra* | 1 | 8 | 29.7 | 237.6 |
| *Paradisaea decora* | 2 | 6 | 28.6 | 171.6 |
| *Paradisaea guilielmi* | 1 | 3 | 4.1 | 12.3 |
| *Paradisaea rudolphi* | 1 | 2 | 58.1 | 116.3 |
| *Cicinnurus magnificus* | 3 | 18 | 146.4 | 2634.4 |
| *Cicinnurus respublica* | 2 | 4 | 36.8 | 147.3 |
| *Cicinnurus regius* | 2 | 3 | 72.3 | 216.9 |
| *Astrapia mayeri* | 1 | 1 | 26.2 | 26.2 |
| *Astrapia rothschildi* | 2 | 3 | 15.5 | 46.6 |
| *Astrapia splendidissima* | 1 | 4 | 53.4 | 213.7 |
| *Astrapia nigra* | 1 | 2 | 50.9 | 101.8 |
| *Epimachus meyeri* | 2 | 5 | 80.8 | 403.9 |
| *Epimachus fastosus* | 4 | 19 | 278.8 | 5297.1 |
| *Ptiloris magnificus* | 3 | 6 | 155.7 | 934.2 |
| *Ptiloris victoriae* | 1 | 2 | 633.5 | 1267.0 |
| *Ptiloris paradiseus* | 1 | 1 | 3.3 | 3.3 |
| *Lophorina superba* | 1 | 3 | 269.4 | 808.3 |
| *Semioptera wallacii* | 2 | 11 | 39.8 | 438.3 |
| *Drepanornis bruijnii* | 1 | 1 | 147.0 | 147.0 |
| *Drepanornis albertisi* | 1 | 4 | 304.7 | 1218.8 |
| *Seleucidis melanoleucus* | 1 | 3 | 60.2 | 180.5 |
| *Parotia wahnesi* | 6 | 9 | 370.6 | 3335.6 |
| *Parotia sefilata* | 2 | 20 | 140.7 | 2813.2 |
| *Parotia lawesii* | 2 | 5 | 79.9 | 399.6 |
| *Parotia helenae* | 1 | 1 | 52.1 | 52.1 |
| *Parotia carolae* | 7 | 11 | 200.6 | 2207.0 |
| *Pteridophora alberti* | 1 | 2 | 76.6 | 153.2 |
| *Manucodia comrii* | 1 | 3 | 9.6 | 28.7 |
| *Phonygammus keraudrenii* | 1 | 1 | 80.2 | 80.2 |

**Supplementary Table 7.** Partial summary (PC1-PC3) of principal components analysis of 5739 notes produced by 32 BOP species. PC loadings for PC1-PC3 were used to plot notes in 3-dimensional PCA space prior to agglomerative hierarchical clustering based on Euclidean distances to categorize notes.

|  | PC1 | PC2 | PC3 |
| --- | --- | --- | --- |
| Summary statistic |  |  |  |
| SD | 2.7348 | 1.6567 | 1.3759 |
| Proportion variance | 0.4986 | 0.183 | 0.1262 |
| Cumulate proportion of variance explained | 0.4986 | 0.6816 | 0.8078 |
| Variable | *loadings* |  |  |
| Duration (sec) | 0.07 | -0.21 | **0.64** |
| Robust Duration (sec) | 0.09 | -0.20 | **0.63** |
| <sup>1</sup>Average entropy | **0.33** | -0.10 | -0.17 |
| <sup>2</sup>Aggregate entropy | **0.33** | -0.11 | -0.14 |
| Min Entropy (bits) | **0.31** | -0.04 | -0.19 |
| Max Entropy (bits) | **0.32** | -0.12 | -0.05 |
| Bandwidth (Hz) | **0.33** | -0.14 | -0.10 |
| Peak Frequency (Hz) | 0.24 | **0.38** | 0.10 |
| Min Frequency (Hz) | 0.12 | **0.53** | 0.19 |
| Max Freq (Hz) | **0.34** | 0.12 | -0.01 |
| <sup>3</sup>PFC Average Slope (Hz/ms) | 0.04 | -0.06 | **-0.18** |
| <sup>3</sup>PFC Max Frequency(Hz) | **0.32** | 0.17 | 0.10 |
| <sup>3</sup>PFC Max Slope (Hz/ms) | **0.27** | -0.24 | 0.03 |
| <sup>3</sup>PFC Min Freq (Hz) | 0.12 | **0.54** | 0.12 |
| <sup>3</sup>PFC Min Slope (Hz/ms) | **-0.27** | 0.20 | -0.08 |

<sup>1</sup> Average entropy describes the amount of disorder for a typical spectrum within the selection by measuring the average of the entropy values for each time slice of the spectrogram.
<sup>2</sup> Aggregate entropy measures the overall disorder in the sound by measuring the energy distribution in each frequency bin of the spectrogram.
<sup>3</sup>PFC = Peak frequency contour

**Supplementary Table 8.** Multiple phylogenetic least-squares (mPGLS) analyses of communication-relevant influences on three axes of courtship phenotype richness conducted only on species without imputed species level values.

| Response variable | Predictor variable | Value | Std.Error | t-value | p-value |
| --- | --- | --- | --- | --- | --- |
| Color richness (log) | (Intercept) | 3.11 | 0.63 | 4.91 | 0.000 |
|  | Behavioral richness (log) | 0.07 | 0.24 | 0.28 | 0.779 |
|  | Acoustic richness (log) | 0.18 | 0.21 | 0.86 | 0.400 |
|  | Display height | 0.28 | 0.14 | 2.01 | *0.055 |
|  | Display proximity | 0.22 | 0.12 | 1.75 | *0.092 |
| Behavioral richness (log) | (Intercept) | 2.12 | 0.58 | 3.67 | 0.001 |
|  | Color richness (log) | 0.04 | 0.16 | 0.28 | 0.779 |
|  | Acoustic richness (log) | 0.38 | 0.15 | 2.49 | **\*0.019** |
|  | Display height | -0.33 | 0.11 | -3.10 | **\*0.004** |
|  | Display proximity | 0.19 | 0.10 | 1.90 | *0.068 |
| Acoustic richness (log) | (Intercept) | -0.80 | 0.79 | -1.01 | 0.320 |
|  | Behavioral richness (log) | 0.49 | 0.20 | 2.49 | **\*0.019** |
|  | Color richness (log) | 0.15 | 0.17 | 0.86 | 0.400 |
|  | Display height | 0.28 | 0.13 | 2.22 | **\*0.035** |
|  | Display proximity | -0.18 | 0.12 | -1.51 | 0.143 |

\*Indicates significant relationships in the ‘full’ analyses incorporating imputed character values.

**Supplementary Table 9.**
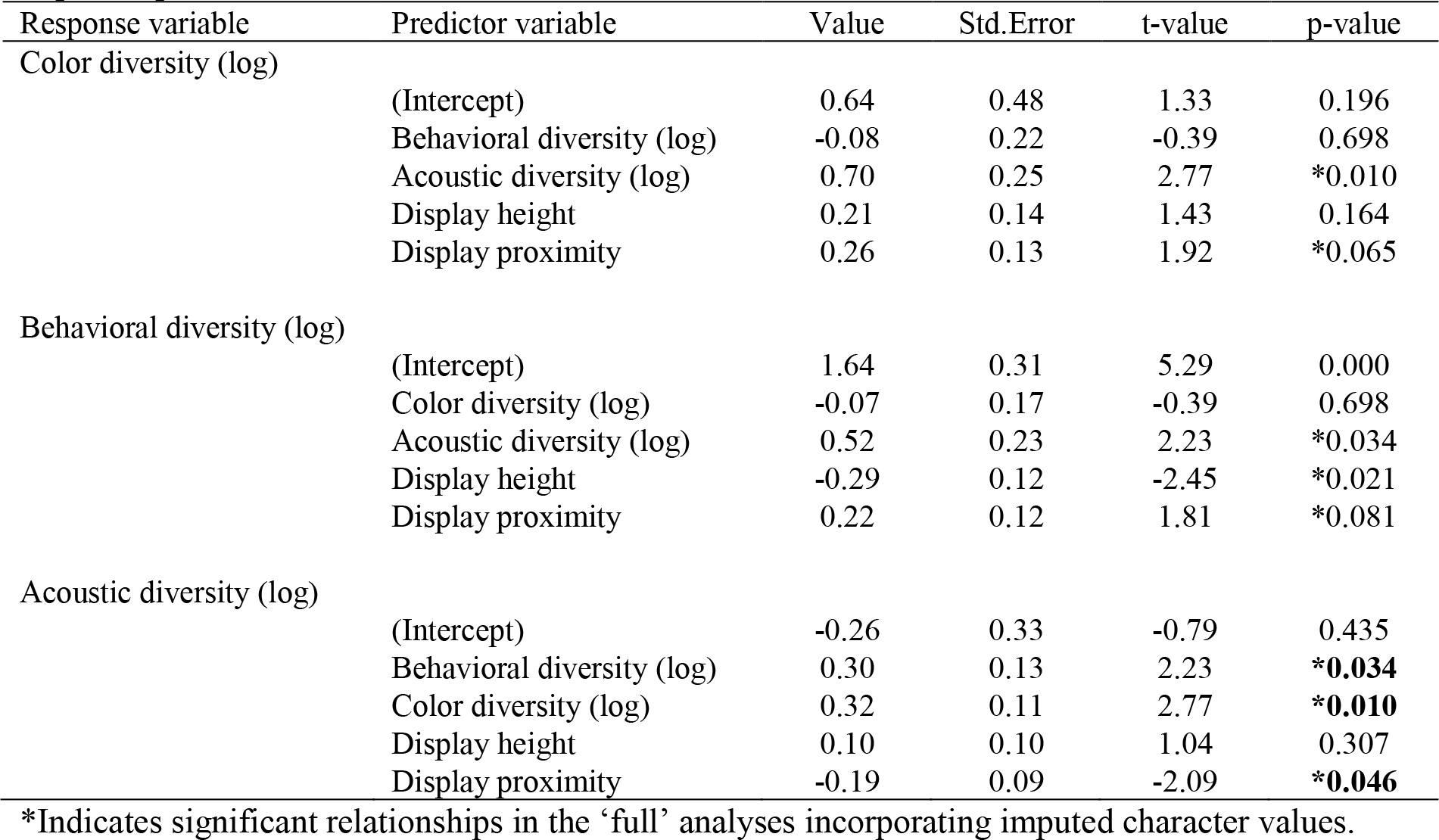
Multiple phylogenetic least-squares (mPGLS) analyses of communication-relevant influences on three axes of courtship phenotype diversity conducted only on species without imputed species level values.

| Response variable | Predictor variable | Value | Std.Error | t-value | p-value |
| --- | --- | --- | --- | --- | --- |
| Color diversity (log) | (Intercept) | 0.64 | 0.48 | 1.33 | 0.196 |
|  | Behavioral diversity (log) | -0.08 | 0.22 | -0.39 | 0.698 |
|  | Acoustic diversity (log) | 0.70 | 0.25 | 2.77 | *0.010 |
|  | Display height | 0.21 | 0.14 | 1.43 | 0.164 |
|  | Display proximity | 0.26 | 0.13 | 1.92 | *0.065 |
| Behavioral diversity (log) | (Intercept) | 1.64 | 0.31 | 5.29 | 0.000 |
|  | Color diversity (log) | -0.07 | 0.17 | -0.39 | 0.698 |
|  | Acoustic diversity (log) | 0.52 | 0.23 | 2.23 | *0.034 |
|  | Display height | -0.29 | 0.12 | -2.45 | *0.021 |
|  | Display proximity | 0.22 | 0.12 | 1.81 | *0.081 |
| Acoustic diversity (log) | (Intercept) | -0.26 | 0.33 | -0.79 | 0.435 |
|  | Behavioral diversity (log) | 0.30 | 0.13 | 2.23 | **\*0.034** |
|  | Color diversity (log) | 0.32 | 0.11 | 2.77 | **\*0.010** |
|  | Display height | 0.10 | 0.10 | 1.04 | 0.307 |
|  | Display proximity | -0.19 | 0.09 | -2.09 | **\*0.046** |

\*Indicates significant relationships in the ‘full’ analyses incorporating imputed character values.

**Supplementary Figure 1.**
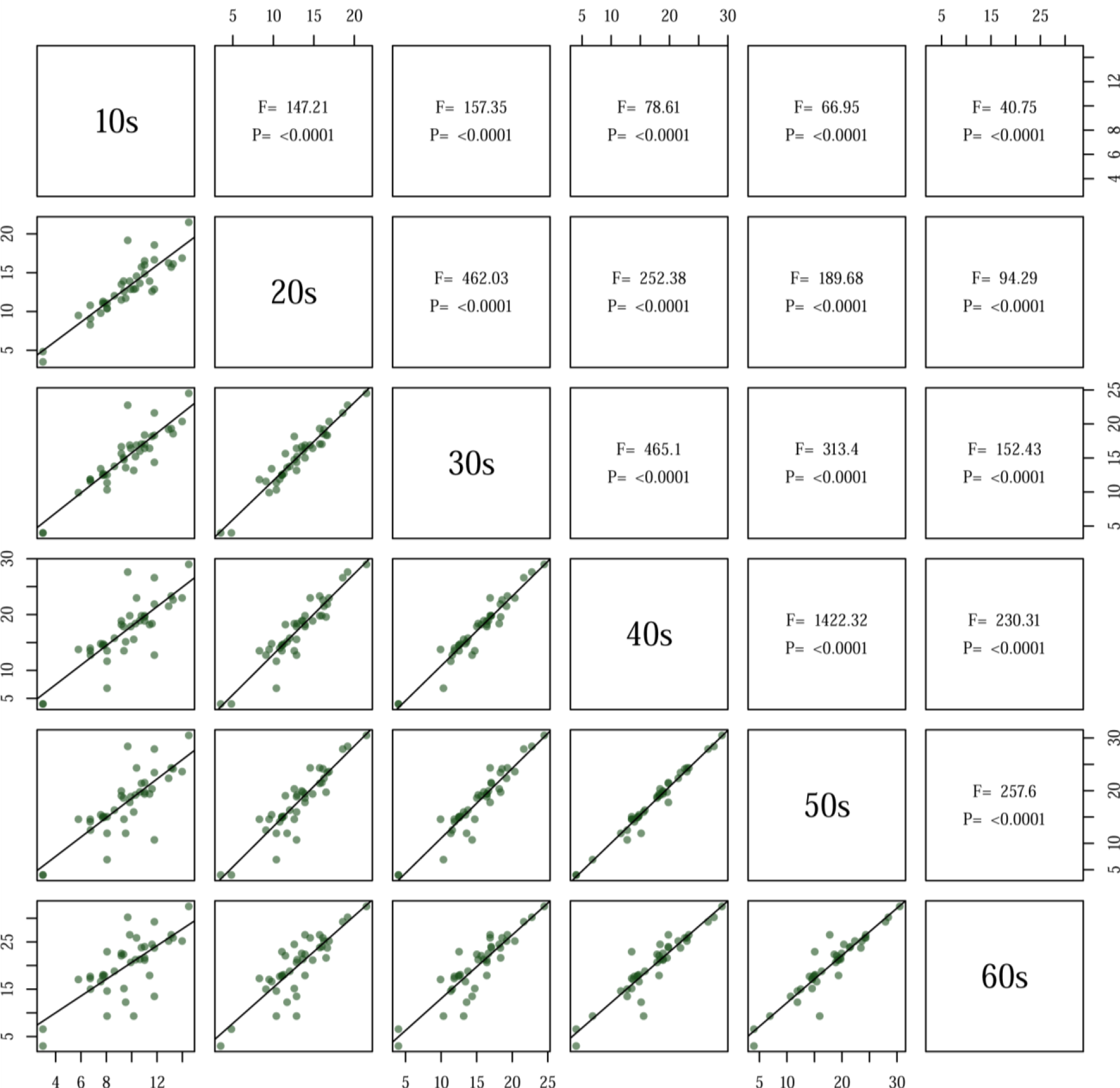
Pairwise comparisons of behavioral richness (number of unique behaviors) estimates for windows between 10 and 60 seconds in duration. Within plots, each point represents a species in the family Paradisaeidae, with species-specific values obtained from rphylopars reconstructions incorporating intra- and interspecific variation. Best-fit lines in lower plots, as well as F and P values presented in corresponding upper diagonal squares, come from PGLS (phylogenetic generalized least squares) analysis assuming Ornstein-Uhlenbeck error structure. Results are qualitatively identical assuming different correlation structures (e.g. Pagel, Brownian).

**Supplementary Figure 2.**
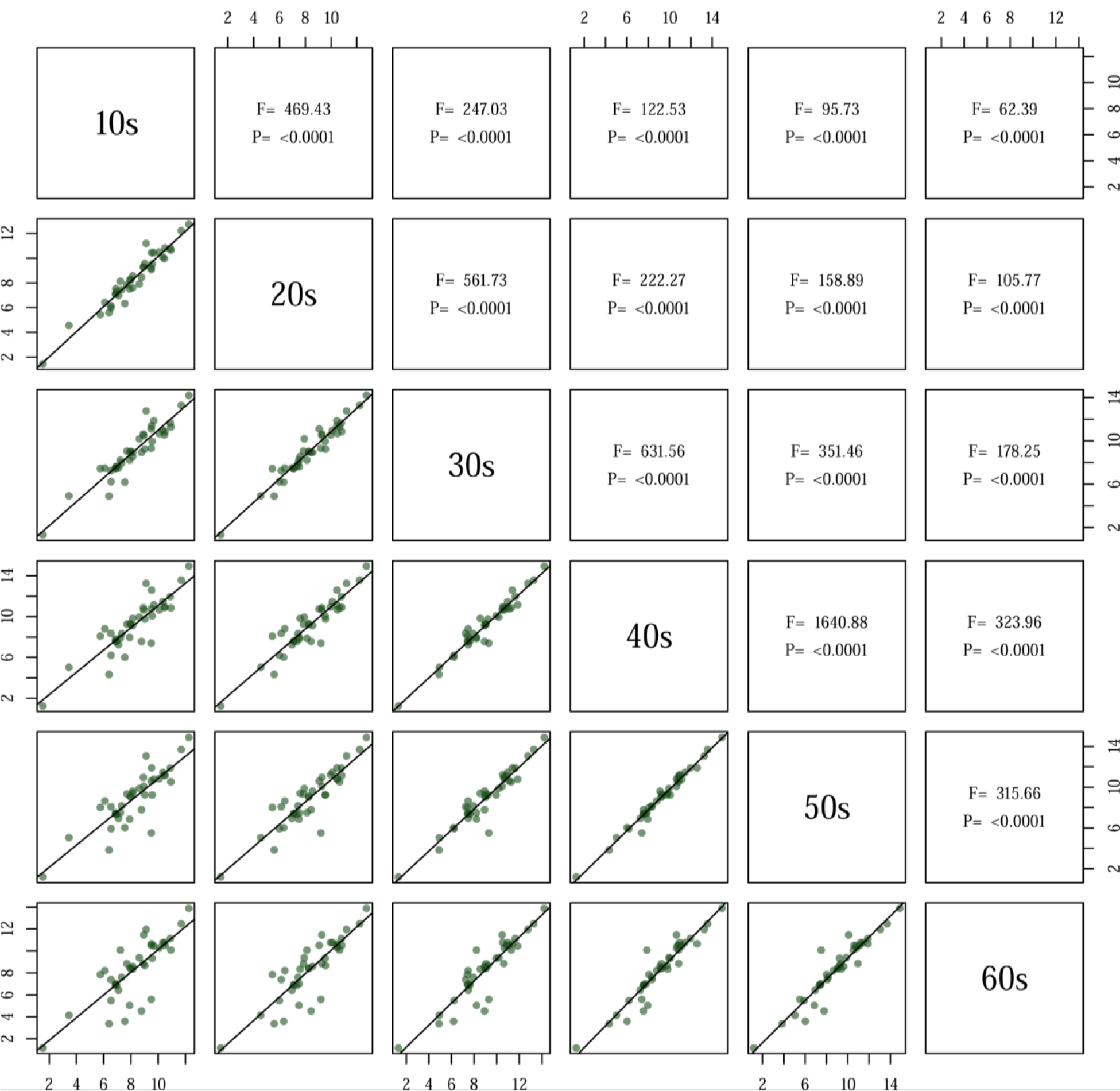
Pairwise comparisons of behavioral diversity (Shannon indices of behaviors) estimates for windows between 10 and 60 seconds in duration. Within plots, each point represents a species in the family Paradisaeidae, with species-specific values obtained from rphylopars reconstructions incorporating intra- and interspecific variation. Best-fit lines in lower plots, as well as F and P values presented in corresponding upper diagonal squares, come from PGLS (phylogenetic generalized least squares) analysis assuming Ornstein-Uhlenbeck error structure. Results are qualitatively identical assuming different correlation structures (e.g. Pagel, Brownian).

**Supplementary Figure 3.**
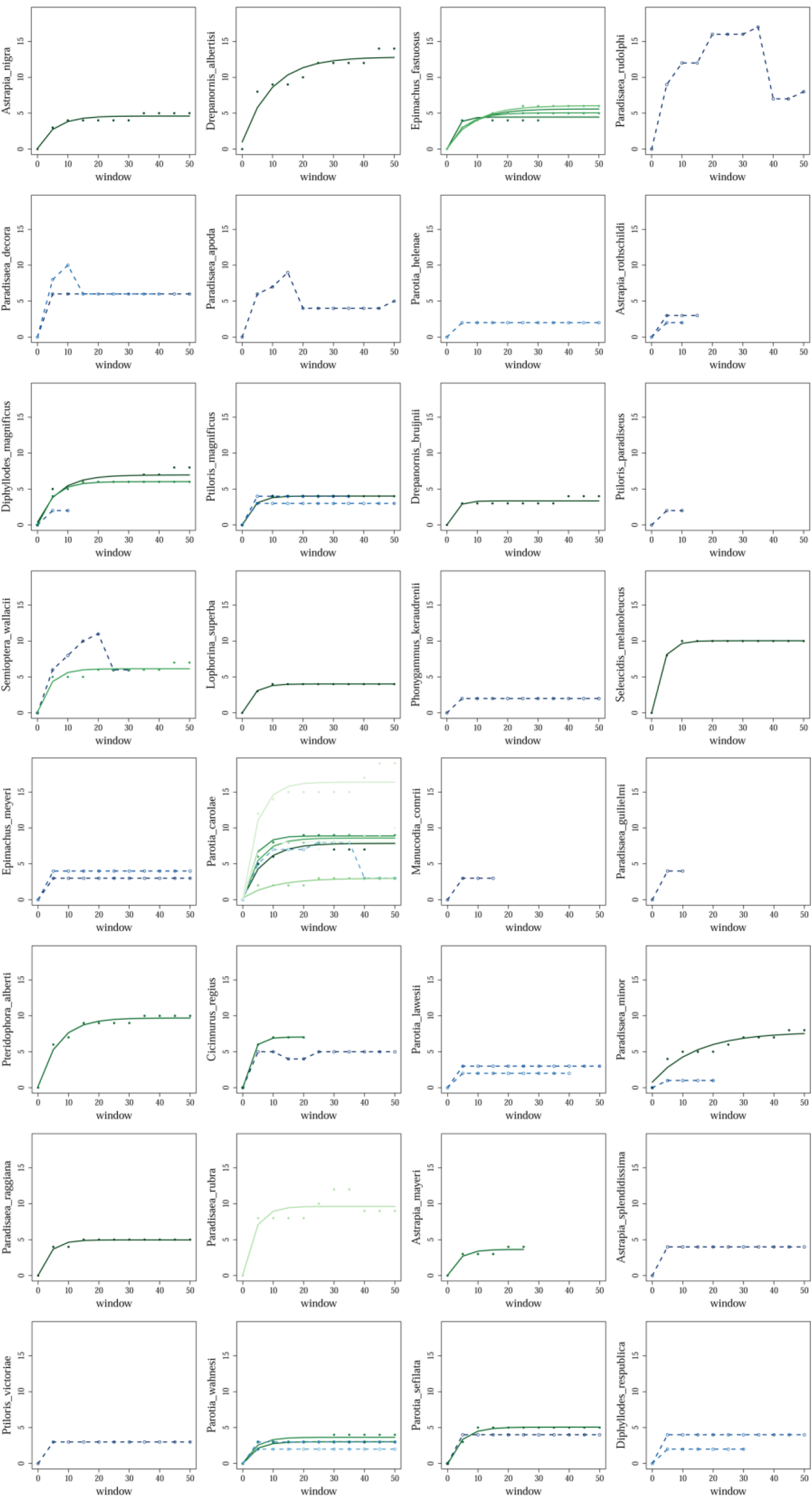
Accumulation curves showing that unique sounds do not continue to accumulate with longer time windows analyzed, across species.

**Supplementary Figure 4.**
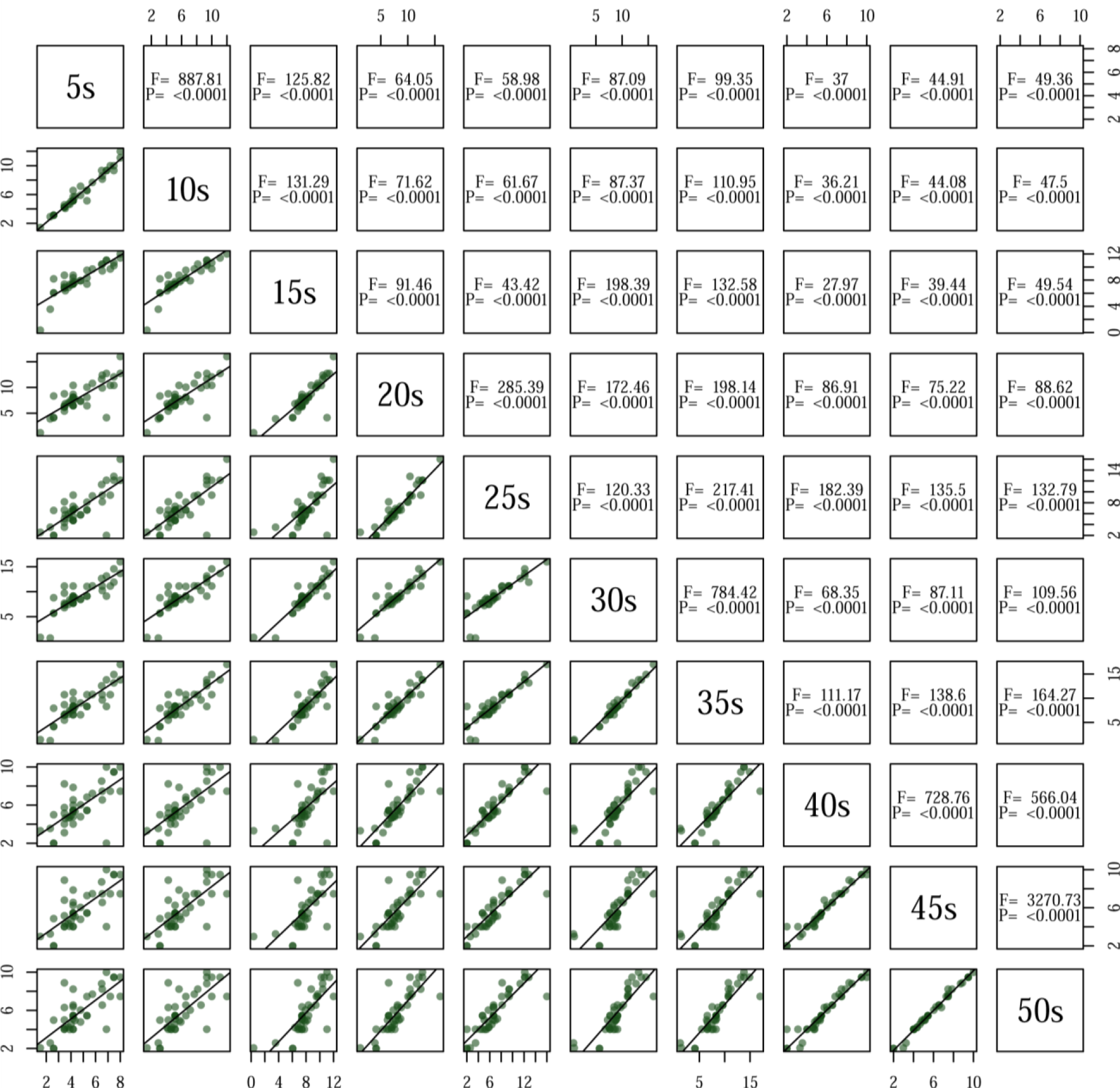
Pairwise comparisons of acoustic richness (number of unique note types) estimates for windows between 5 and 50 seconds in duration. Within plots, each point represents a species in the family Paradisaeidae, with species-specific values obtained from rphylopars reconstructions incorporating intra- and interspecific variation. Best-fit lines in lower plots, as well as F and P values presented in corresponding upper diagonal squares, come from PGLS (phylogenetic generalized least squares) analysis assuming Ornstein-Uhlenbeck error structure. Results are qualitatively identical assuming different correlation structures (e.g. Pagel, Brownian).

**Supplementary Figure 5.**
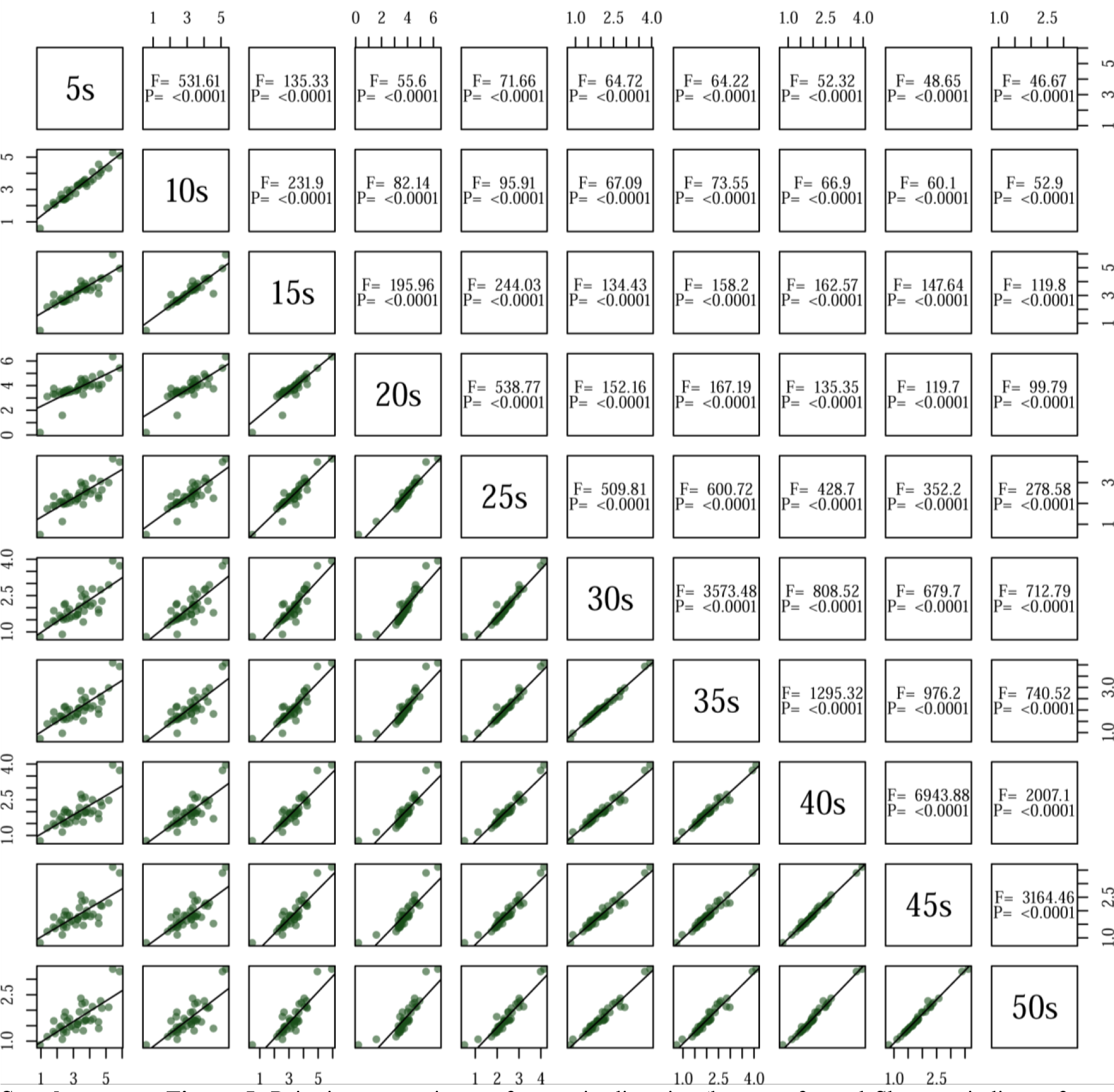
Pairwise comparisons of acoustic diversity (log transformed Shannon indices of note complexity) estimates for windows between 5 and 50 seconds in duration. Within plots, each point represents a species in the family Paradisaeidae, with species-specific values obtained from rphylopars reconstructions incorporating intra- and interspecific variation. Best-fit lines in lower plots, as well as F and P values presented in corresponding upper-diagonal squares, come from PGLS (phylogenetic generalized least squares) analysis assuming Ornstein-Uhlenbeck error structure. Results are qualitatively identical assuming different correlation structures (e.g. Pagel, Brownian).

